# Evolution to increased positive charge on the viral spike protein may be part of the adaptation of SARS-CoV-2 to human transmission

**DOI:** 10.1101/2022.07.30.502143

**Authors:** Matthew Cotten, My V.T. Phan

## Abstract

The severe acute respiratory syndrome coronavirus 2 (SARS-CoV-2), the causative agent of the coronavirus disease 2019 (COVID-19) pandemic, continues to evolve and infect individuals. The exterior surface of the SARS-CoV-2 virion is dominated by the spike protein and the current work examined spike protein biochemical features that have changed during the 2 years that SARS-CoV-2 has infected humans. These biochemical properties may influence virion survival and promote movement through the environment and within the human airway to reach target cells to bind, enter and establish the next round of infection. In addition to selective pressure to avoid immune recognition of viral proteins, we hypothesised that SARS-CoV-2 emerged from an animal reservoir capable of human infection and transmission but in a sub-optimum state and a second level of selective pressure is acting on these biochemical features. Our analysis identified a striking change in spike protein charge, from −8.3 in the original Lineage A and B viruses to −1.26 in the current Omicron viruses. In summary, we conclude that in addition to immune selection pressure, the evolution of SARS-CoV-2 has also altered viral spike protein biochemical properties. Future vaccine and therapeutic development should also exploit and target these biochemical properties.

## Introduction

The severe acute respiratory syndrome coronavirus 2 (SARS-CoV-2), the causative agent of the coronavirus disease 2019 (COVID-19) epidemic, continues to evolve and infect individuals. Similar to other viruses, the SARS-CoV-2 virion biochemical properties play an important role in controlling virus transmission. After replication in an infected individual and release from an infected cell, onward transmission requires survival of the virion to reach susceptible cells in a new host individual initiating the next round of infection. The physical properties of the surface proteins of the virus such as charge, size, hydrophobicity and folding may influence movement of the virion through the environment, promoting or limiting binding of the virion to the external surfaces. Once reaching a susceptible individual, virion physical properties may influence movement within the human airway and determine the ability of an infecting virion to reach target cells to bind, enter and replicate (Adamczyk et al. 2021). The exterior surface of the SARS-CoV-2 virion is dominated by the spike protein and the current work examines simple spike protein features that have changed during the 2 years of the SARS-CoV-2 pandemic. In addition to selective pressure to avoid immune recognition of viral proteins, we hypothesise that SARS-CoV-2 emerged from an animal reservoir capable of human infection and transmission but in a sub-optimum state. Additionally, there is a second level of selective pressure to adjust to the physical transmission between humans. Evidence for this adaptation can be found in changes in the SARS-CoV-2 spike protein over recent evolution. With over 11 million SARS-CoV-2 genomic sequences generated globally from across the pandemic, many of these sequences have intact spike gene sequences that can be used to monitor change across the 2 years of human host evolution of this virus.

Much of the observed spike protein substitutions may be in response to the developing immune response to this new pathogen, which is reflected in substitutions occurring in the immune-exposed S1 domain of the spike protein and there is ample evidence that many of these spike protein changes allow escape from host immunity (Tzou et al. 2022)(Greaney, Loes, et al. 2021)(Greaney, Starr, et al. 2021)(Greaney et al. 2022) (Cao et al. 2022) (Dejnirattisai et al. 2022) (DeGrace et al. 2022). There may also be evolutionary selection for protein changes that improve host interactions apart from immune evasion. These include altering spike/receptor binding kinetics, protease cleavage events, tertiary structure (S1/S2 interactions after cleavage) or the physical properties of the virion (charge, hydrophobicity, and protein folding or secondary structure) in ways that might improve transmission. To explore the role of the biochemical features of the spike protein in human transmission, we monitored changes in spike biochemical features over the two years that SARS-CoV-2 has been evolving in humans and report an increase in spike protein positively charge especially among the virus lineages that were highly prevalent.

## Results

The SARS-CoV-2 spike protein physical features were calculated from spike protein sequences from across 2 years of the COVID-19 epidemic. Features that could be quantitated from protein sequence were used (see Methods), including charge at pH 7.4, Kyle and Doolittle GRAVY score (Kyte & Doolittle 1982) (which is a measure of hydrophobicity), an instability index derived from dipeptide content (Guruprasad et al. 1990), properties influencing protein folding (percent helix, fold or sheet as predicted from amino acid content), individual amino acid total fraction and di-amino acid total fraction.

A dominant pattern of SARS-CoV-2 evolution during the two years of human adaptation has been the regular appearance and the subsequent regional and then global dominance of lineages. These lineages typically encode a small set of amino acid changes from earlier lineages, many of which are likely to provide temporary or long-term advantage for the viral lineage. An analysis was performed to identify spike physical features most strongly linked with SARS-CoV-2 lineages (Figure 1). The first 300 reported genomes from each major lineage were collected, spike protein sequences were extracted and the physical features of each protein were collected into a matrix. The top features distinguishing SARS-CoV-2 lineages were identified with charge as the most important feature (Figure 1A). A principal component analysis using the top 8 features (charge, gravy, fraction T, fraction R, instability, fraction G, fraction D and fraction K), provided clustering of spike sequences by lineage (Figure 1B). These results support the idea that spike protein charge (among other features) is an important determinant of the lineages that have evolved during the first two years of the COVID-19 epidemic.

**Figure 1.**
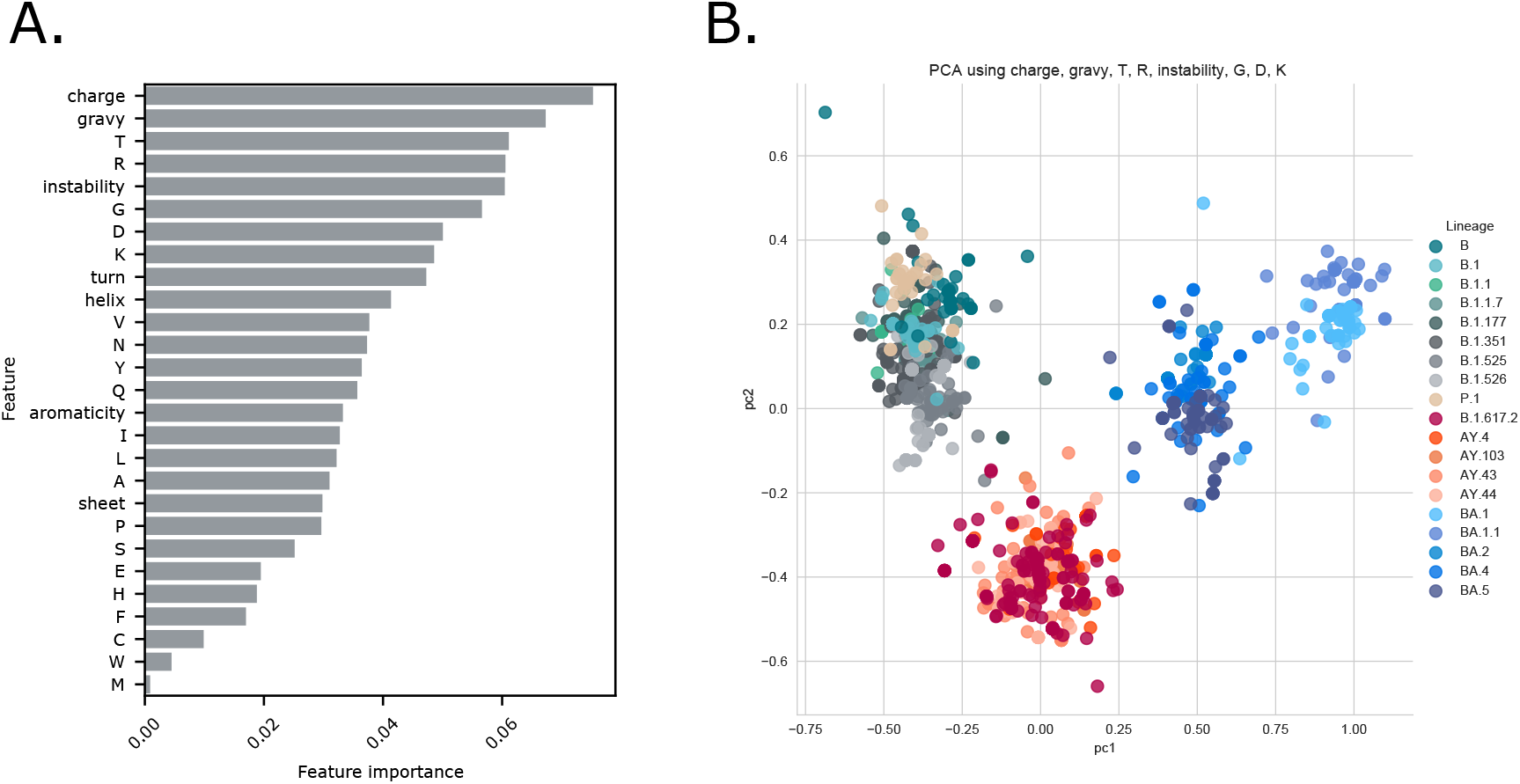
Identification of spike protein charge association with SARS-CoV-2 lineage. **Panel A:** A set of 300 spikes sequences extracted from the first 300 SARS-CoV-2 genomes per lineage (by date of collection) was analyzed, features for each sequence were collected (see Methods). SKLearn feature selection (Pedregosa, F. and Varoquaux, G. and Gramfort, A. and Michel, V. et al. 2011) was used to identify features that most accurately identified the sequence lineage. The importance of features were ranked in order. **Panel B:** The top 8 features (charge, gravy, fraction T, fraction R, instability, fraction G, fraction D, fraction K) were further used in a principal component analysis to cluster the same set of SARS-CoV-2 spike sequences. Each node represents a single spike sequence, nodes were coloured by Pangolin lineage assigned to the genome from which the spike sequence was obtained. Lineage colouring is explained in the figure legend to the right.

Changes in charge of spike protein across the epidemic were investigated. Plotting total spike charge for all genomes per month of the epidemic showed a clear pattern of increase in charge over two years of evolution (Figure 2, panel A). Median spike charge was −8.3 in the original SARS-CoV-2 viruses reported in late 2019 to early 2020, by March 2020, an increase in positive charge to −7.28 was observed. Subsequently, an additional increase in positive charge occurred in mid-2021 to −3.28, and most recently a charge increase occurred in late 2020/early 2021 to −1.26.

**Figure 2.**
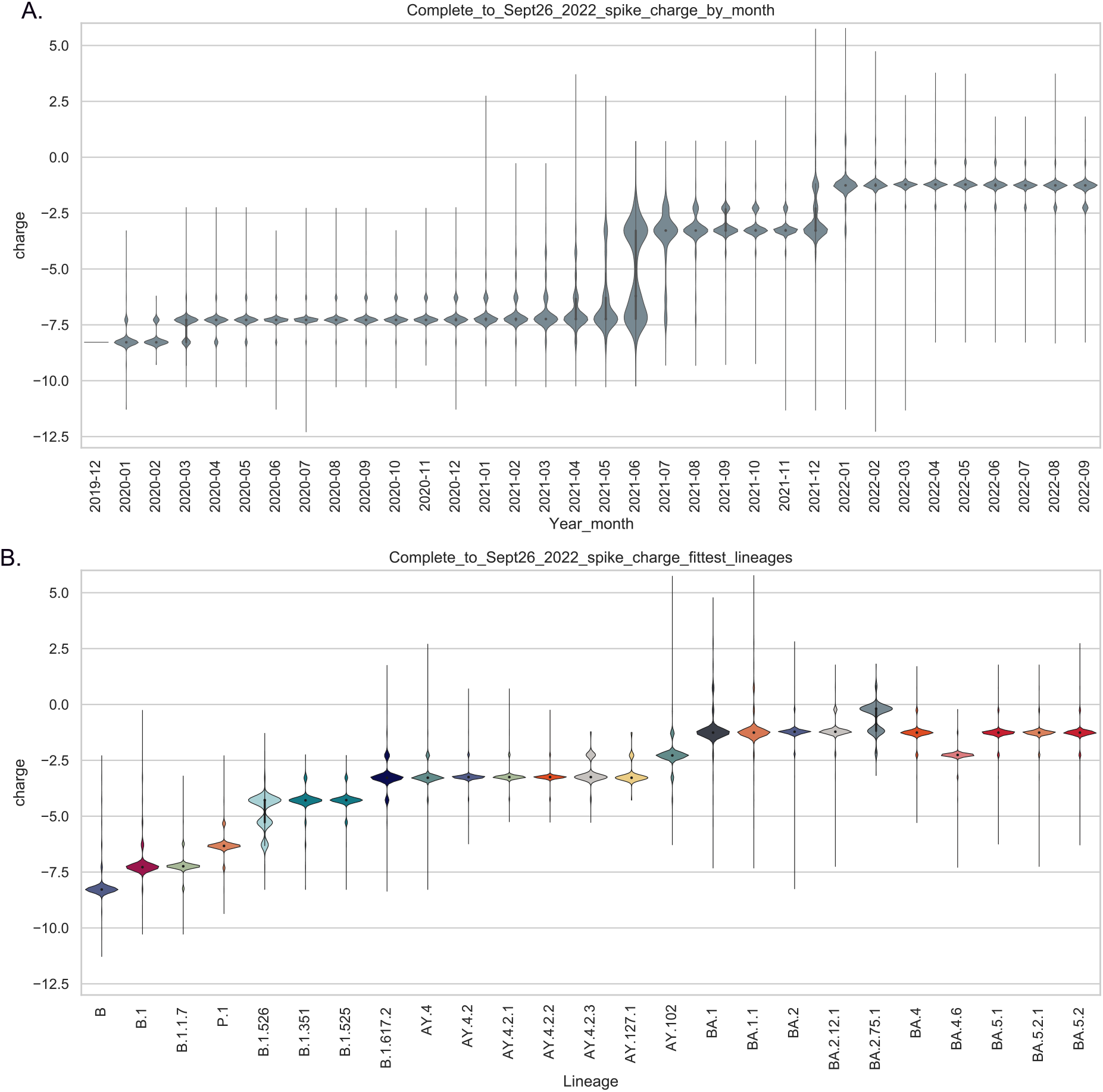
**Panel A. Total SARS-CoV-2 spike charge per epidemic month.** All available SARS-CoV-2 genomes up to 24 September 2022 were retrieved from GISAID (GISAID 2020) and the spike protein sequence was extracted (if intact). Total charge at pH 7.4 was calculated and values were plotted using a violin plot by month of sample collection. For each epidemic month the violin plot depicts the distributions of calculated spike charge for all available SARS-CoV-2 genomes. **Panel B. Spike charge in major SARS-CoV-2 lineages**. For each lineage, all available spike sequences were collected (up to 24 September 2022), total charge was measured and violin plots prepared to show the charge distribution by lineage. Lineages (indicated at bottom of chart) were ordered by their appearance in the epidemic.

These spike protein charge increases can be attributed to the major successful lineages reported over time (Figure 2B). The B.1, B.1.1 and B.1.1.7 (Alpha) lineages that dominated the first year of the epidemic encoded spike proteins with charges between −8 and −6 while the B.1.351 (Beta) and B.1.525 (Eta) lineages showed a further increase in charge to around −4.5. The B.1.617.2 (Delta) lineage and sublineages (AY.x) displayed further increase in charge. Most recently, the Omicron variants (including BA.1, BA.1.1, BA.2, BA.2.12,BA.3, BA.4 and BA.5) show further spike charge increases with the majority of Omicron encoded spike proteins showing charge at −1.26 (Figure 2, panel B).

Some indication of functional consequences of the observed changes in spike charge can be obtained from the location on the charged amino acid substitutions in the spike protein. Sets of spike sequences (extracted from the first 300 reported genomes per select lineage) were processed to illustrate the changes to more negative charge (blue) or more positive charge (orange/red) in the protein relative to the initial Lineage B genome sequences (Supplementary Figure 1). The initial change in charge was a substitution of an aspartic acid residue (D, with a calculated charge of −1) by a glycine (G, neutral). In some early lineages (e.g. A.23.1), proline (P) at position 681was substituted with the positively charged arginine (R),or Q680 was substituted with a partially charged histidine H residue. The P681R positive substitution promotes furin cleavage and activation of the spike protein for cell fusion (Lubinski et al. 2022)(Liu et al. 2022). The Delta lineage spike proteins encoded additional positive charge in the ACE2 binding region, as well as in the far amino terminal region and near the heptad repeat (HR1) which may also enhance membrane fusion activity. More recently, a number of positive substitutions have occurred in the Omicron lineage virus spike proteins with predominance of positively charged changes in the receptor binding domain (Supplementary Figure 1), suggesting a role of increased charge in spike/receptor interactions.

It is probable that the spike protein has an upper limit to the amino acid charge that it can allow for proper folding, assembly and function. This upper charge value will be determined by the acquisition of optimum transmission properties in balance with immune selection. After the regular increase of spike protein charge observed up to the appearance of the Omicron lineages, an indication of a stasis in positively-charged amino acid accumulation is now displayed by SARS-CoV-2 Omicron lineages. The majority of Omicron sub-lineages remain at spike charge −1.26 (Figure 3A) although a few specific Omicron sub-lineages show changes toward more positive or negative charge (e.g. BA.2.75.1 more positive, BA.4.6 more negative, as illustrated in Figure 2B) with the additional changes often associated with immune selection. To monitor the current trends of spike protein changes, we calculated the fraction of reported genomes with spike charge greater than or less than the Omicron mean charge of −1.26 and documented how these fractions had changed over the last 4 months of the pandemic (Figure 3B). The majority of encoded spike proteins are almost exclusively from Omicron lineage viruses and show a charge of −1.26. However, a small fraction of genomes encode spikes proteins with slightly more or less charge (Figure 3B) with the greater trend (20% of all reported genomes in September 2022) showing more negative charge (Figure 3B).

**Figure 3.**
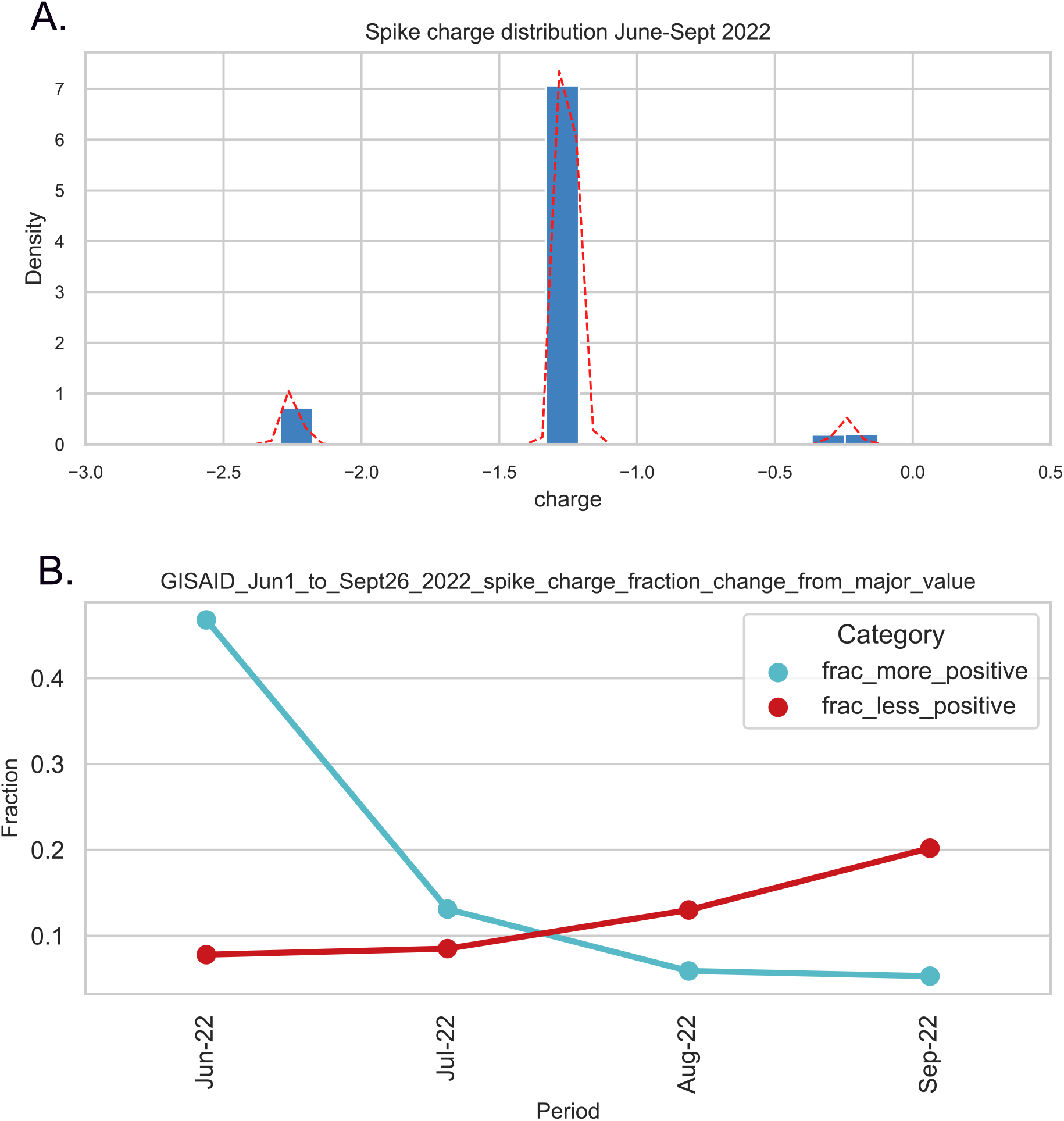
Recent changes in spike protein charge. **Panel A**: All available spike proteins from genomes with sample collection dates of June-Sept 2022 were analyzed for total spike charge. A histogram of the calculate total spike charges for the entire set is shown in panel A with the **kernel density estimation** (**KDE**) line in red. A major peak at −1.26 is observed wit small outlier peaks of genomes with more negative and more positive spike proteins **Panel B**: For each month (over the period June 1 to Sept 26 2022) the fraction of reported genomes for that month with charge greater than or less than the majority value of −1.26 was calculated.

Lastly, we investigated if a similar pattern of spike charge evolution could be observed in other coronaviruses that have made a transition to human transmission. In recent history, several coronaviruses (in addition to SARS-CoV-2) have been observed to jump hosts. For example, coronavirus 229E is commonly detected in humans and very close coronaviruses have been identified in bats (Victor Max Corman et al. 2015) (Tao et al. 2017) and camels (Victor M. Corman et al. 2016) (Sabir et al. 2016) suggesting movement of the virus between hosts. All available coronavirus 229E full genomes sequences were retrieved from GenBank, the spike coding region was extracted from the genomes, translated and total charge was calculated. A difference from −26 to −8, or almost 18 charge units is seen comparing 229E-like viruses from bats to 229E from humans (Figure 4A) and almost 9 charge unit difference was observed in spike median charge comparing 229E viruses from camel vs. human infections (Figure 4A).

**Figure 4.**
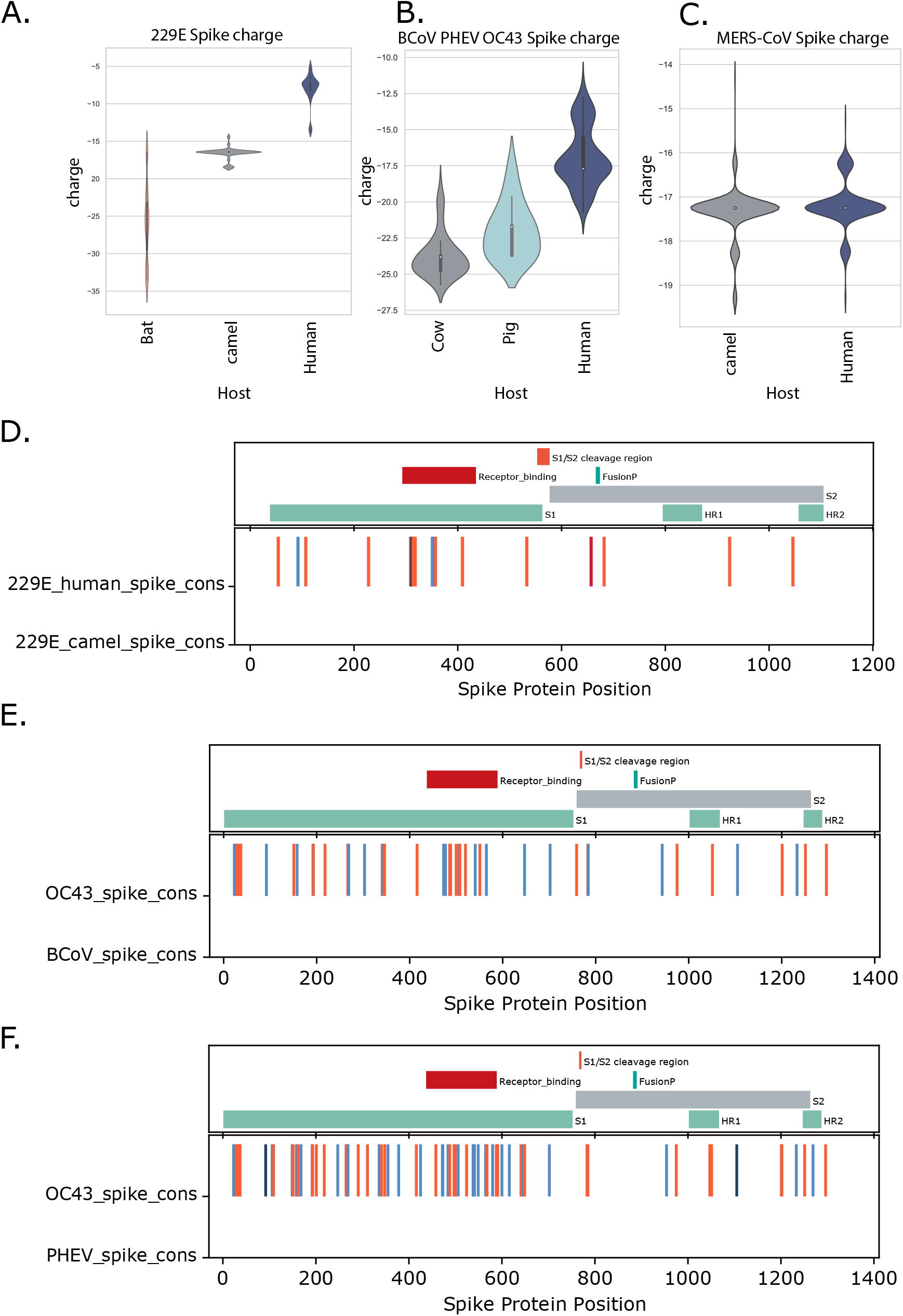
**Spike charges from select groups of coronaviruses that have moved into humans (Panels A-C).** All available full genomes for the indicated coronaviruses were retrieved from GenBank, the spike coding region was identified and translated into protein and total charge at ph 7.4 was calculated. Violin plots indicate the charges of each collection of spike proteins, median values are indicated by the open square. **Panel A**. Coronavirus 229E from bat, camel or human infections, **Panel B**. BCoV (from bovine infections) PHEV (from porcine infections) and OC43 (from human infection), **Panel C**. MERS-CoV from camel or human infection. **Panel D-F: Consensus spike protein sequences were generated from the indicated virus groups and charged amino acid changes were determined**. Charge changes were colored from dark blue (change from positively to negatively charged amino acid (AA)), blue change from neutral to negatively charged AA), orange (change from neutral to positively charged AA) and red (change from negative to positively charged AA). **Panel D:** 229E spike from human infections compared to 229E spike from camel infections, **Panel E**: Human OC43 spike compared to BCoV spike, **Panel F**: Human OC43 spike compared to PHEV spike. Key spike protein features of each group’s spike protein are shown in the upper portion of each panel.

Infection with coronavirus OC43 is common in humans and closely related viruses are found in cattle (bovine coronavirus, BCoV) (Vijgen et al. 2006) and pigs (porcine hemagglutinating encephalomyelitis virus, PHEV) (Vijgen et al. 2005) (Vijgen et al. 2006). Comparing the three OC43-type virus groups, the human virus OC43 has an increased charge of ca. 5 units compared to PHEV and ca.7 units compared to BCoV (Figure 4B).

The commonly known host for the Middle East Respiratory Syndrome coronavirus (MERS-CoV) is dromedary camels; however, zoonosis and serious human infections occur frequently (Cotten et al. 2013) (Cotten et al. 2014) (Memish et al. 2014) (Zhou et al. 2021) (So et al. 2019) as reviewed in (Peiris & Perlman 2022). From 698 full MERS-CoV genomes available in GenBank, there was no strong difference in the encoded spike charge of virus sequences derived from human versus camels infections (Figure 4C).

Considering the location of the charge differences in the spike proteins, for coronavirus 229E, the charge increases occurred throughout the protein, although there is a slightly higher number of positive changes in the receptor binding region of the human infection derived viruses (Figure 4D). For OC43, the porcine and human viruses also show increases in positive charge throughout the spike protein, the porcine PHEV also showed a slight enrichment in positive charge in the receptor binding region (Figure 4E).

## Discussion

After more than two years of the COVID-19 pandemic and with the availability of >11 million SARS-CoV-2 genome sequences, a trend of SARS-CoV-2 spike protein charge can be observed, with successive lineages showing an increase in positive charge over earlier lineages. Over the course of the pandemic, the SARS-CoV-2 spike protein has evolved from a protein with a total charge of −8.28 in the original Lineage A and B viruses to a protein with a total charge of −1.26 in the majority of the currently circulating Omicon lineage viruses. This pattern has been noted previously (Pawłowski 2021) (Nie et al. 2022). We expand on these observations, and document lineage patterns and sites of change in the spike protein and explore similar phenomena of evolution to more positive charge in two other coronaviruses (OC43 and 229E) that have moved between animals and humans.

This study does not identify a mechanistic basis for the increased spike charge although there are several possible transmission steps that might be promoted by increasing charge. Exposed, positively charged spike amino acids should promote interactions with negatively charged cellular structures. Interactions with negatively-charged heparin have been reported with SARS-CoV-2 spike (Kim et al. 2020) and negatively-charged sialylated glycans are reported to promote entry of SARS-CoV-2 (Nguyen et al. 2022). The upper respiratory tract is coated by and protected by mucins, frequently modified with sialic acid or phosphorylated, high mannose N-glycans (Byrd-Leotis et al. 2021) which present a negatively charged matrix that could either promote or protect against viral transmission. The SARS-CoV-2, OC43, and BCoV virions display binding to negatively charged carbohydrate structures found in the airway (Byrd-Leotis et al. 2021) and the ionic environment of the human upper respiratory tract may favour binding and transmission of viruses with increased positive charge. Perhaps it is not surprising that both OC43 and 229E coronaviruses exhibited increases in spike positive charge after moving from animal hosts (cow, pig and camel) to human hosts (Figure 5). A similar change in MERS-CoV was not observed, however MERS-CoV currently shows only limited human to human transmission with most known transmission chains ending after 2 to 3 human to human transmission events as shown in (Assiri et al. 2013) and (Cotten et al. 2013). MERS-CoV might not have experienced sufficient number of human replication cycles or have undergone the same level of selection for human transmission that OC43, 229E and SARS-CoV-2 have experienced. For both OC43 and 229E coronaviruses moving to humans, the broad location of the positive changes across the spike protein sequence suggested that positive charge may be promoting several functions including receptor binding, furin cleavage, cell fusion as well as antigenic changes or less specific changes to avoid or promote ionic interactions during transmission.

There is likely a limit to the accumulation of positively charged residues in the SARS-CoV-2 spike protein. Functional constraints exist, there may also be penalties associated with non-specific binding due to excess positive charge, and there are certainly charge influences on protein folding and higher order protein interactions (Creighton 2002). Our prediction is that the SARS-CoV-2 protein will reach some upper limit of charge defined by these constraints. Indeed, we observe that the majority of Omicron lineages encode spike proteins with charge −1.26, after more than 6 months of evolution (Figures 2A and 2B). A small fraction of genomes with more positive charge or less positive charge have appeared, but the global tendency across all reported genomes from June to September 2022 is a modest decline in the positive charge (Figure 3b) which suggest the upper limit to charge has been reached.

Could these changes in spike charge have occurred by chance and not be a response to selective pressure? Of the 20 standard amino acids (AA), only 2 AA have negatively charged side chains, 2 AA have positively charged side chains while the remaining 16 AA are neutral at pH 7.4.. Assuming equal probability of any AA change, there is an 18/20 chance of a negative AA being substituted by a neutral or positively charged AA and the majority of change opportunities would result in loss of negative charge. However, natural selection is more complex, because the genetic code uses 3 adjacent nucleotides to encode an AA, there are multiple encoding possibilities for each AA, the codon redundancy is not identical for each AA and the number of nucleotide changes required to produce any particular AA change can be 1, 2 or 3. This has resulted in an evolved protein stability in the genetic code (Chan et al. 2020) with AA changes that maintain rather than change physical properties (negative, positive, polar, non-polar, aromatic) more likely based on the codon array (Livingstone & Barton 1993) and the nucleotide changes required for an AA change. For example, the probability of a negative AA to negative AA change is 0.333 while the probabilities of change of a negative AA to a non-polar, aromatic, polar or positive AA are 0.051, 0.044, 0.028 and 0.044 respectively, with changes away from a negative charged AA nearly 10-fold less likely to occur than conserving the negative charge at that position (Livingstone & Barton 1993). For these reasons, it appears that the accumulation of positive charge on spike protein has not occurred by chance and is likely providing some selective advantage for the virus. It should also be noted that the observed charge changes in exposed virion proteins seem to be limited to spike. Two additional SARS-CoV-2 proteins are externally exposed, the E protein (ORF4) and the M protein (ORF5), showing no consistent change in the charge of either of these proteins across the 2 years of the epidemic (results not shown).

Obermeyer et al. documented AA substitutions associated with SARS-CoV-2 fitness (Obermeyer et al. 2022). Consistent with the idea that the increase in positive charges is not by chance, of the top 20 substitutions increasing SARS-CoV-2 fitness, 14 substitutions were in the spike protein, among which 4 were changes that increased positive charge while only 1 of 14 introduced a negative charge in spike (Obermeyer et al. 2022).

Natural selection could be acting on multiple features of the spike protein. The necessity to avoid host immune responses is likely to be the major selective force acting on the virus. This results in the amino acid changes, which in turn are determined by epitopes. The selection for increased charge in the spike protein is probably occurring in the background, not as a major shift needed to bypass immune responses. However, the increase in charge may improving survival and transmission in humans in subtle ways, and this advantage, when multiplied over the millions of infections can provide some of the growth and infection advantages seen by new SARS-CoV-2 variants. It is proposed that the N764K, N856K and N969K substitutions (all increasing spike positive charge) may enhance S1/S2 subunit interactions after proteolytic processing of the spike protein, resulting in reduced S1 shedding and improving transmission (Martin et al. 2022) Increased charge may also alter receptor interactions. In the Omicron (BA.1) spike protein, the Q493R and Q498R substitutions are predicted to allow two additional salt bridges with ACE2 receptor position 35Glu and 38Glu (McCallum et al. 2022). Indeed, looking at the timing of charge shifts in each major lineage, the changes to more positive charge accumulate later than the changes that first allow a lineage to emerge and dominate global infections. In this model, the primary spike changes are driven by immune selection and allow a new lineage to bypass existing immune responses. Once a successful new variant emerges, the large number of new infections allow selection for the accumulation of beneficial positive charge changes. The similar pattern of increased positivity of spike protein in other coronaviruses that have moved between animals and humans (OC43, 229E, Figure 4) suggest that the change in surface protein charge may be a more general phenomenon with coronaviruses and might be a useful parameter to examine when monitoring zoonosis. This study provides a framework to monitor viral evolution through changes in biochemical properties, which can be easily applied to other viruses important to public and global health. An important note, our analyses on viral spike protein biochemical properties to monitor virus evolution are not meant to replace traditional phylogenetic analyses. The observed pattern of biochemical properties changes should completement phylogenetic signals. However, in situations where there are limited sequences available to produce reliable phylogenetic signals (e.g. the 229E and OC43 viruses examined in Figure 4), this kind of analysis using virus biochemical properties from different host species would certainly help provide important information on the virus evolution, zoonosis as well as aiding the prediction of patterns of viral changes.

In conclusion, our study provides an novel analytical framework to monitor viral evolution through changes in biochemical properties, which can be easily applied to other viruses important to public and global health. We also showed that natural virus evolution is more complicated and may involve multiple factors including immune selection, as well as spike protein biochemical properties. The observation of increase of SARS-CoV-2 spike protein charge over time provides useful information for future vaccine and therapeutic development.

## Methods

Full alignments of SARS-CoV-2 genomes were obtained from GISAID (GISAID 2020) with collection dates to 15 June 2022. All spaces in fasta IDs were removed using sed (sed -i -e ‘s/ /_/g’ msa_xxxx.fasta), the alignment was dealigned (“-” characters removed) and genomes were classified using Pangolin (Áine O’Toole et al. 2020) with the most recent database updates (pangolin v4.1.1, pangolin-data v1.11 constellations v0.1.10 and scorpio v0.3.17). The spike coding region from each genome (if present and intact (no Ns)) was translated into protein. Features of the protein that could be quantitated from the spike protein sequence were determined using the ProteinAnalysis functions from BioPython (Cock et al. 2009). These features included charge at pH 7.4, Kyle and Doolittle GRAVY score (Kyte & Doolittle 1982) (a measure of hydrophobicity), an instability index derived from dipeptide content (Guruprasad et al. 1990), the total percent helix, fold or sheet properties of the protein and the total fractions of individual amino acids and fractions of di-amino acids. A matrix of all spike protein features plus collection date, and lineage was prepared and used for analysis. Similar analyses were performed for other coronaviruses such as 229E, OC43 and MERS-CoV by retrieving all complete genomes available from GenBank (15 June 2022). The spike protein was also extracted using the same method as aforementioned. Additional details are provided in the figure legends. The python code used for the analyses is available here: https://github.com/mlcotten13/SARS-CoV-2_spike_charge.

## Acknowledgements

We thank all global SARS-CoV-2 sequencing groups for the open sharing of sequence data and to the GISAID platform and team for making these data available. We are grateful to Andrew Rambaut, Áine O’Toole and the Pangolin team for the Pangolin typing tool and resources.

## Funding

This work was supported by the UK Medical Research Council (MRC/UK Research and Innovation) and the UK Department for International Development (DFID) under the MRC/DFID Concordat agreement (grant agreement no. MC_PC_20010) and Wellcome Trust, UK FCDO-Wellcome Epidemic Preparedness-Coronavirus (grant agreement no. 220977/Z/20/Z).

---

**Supplementary Figure 1.**
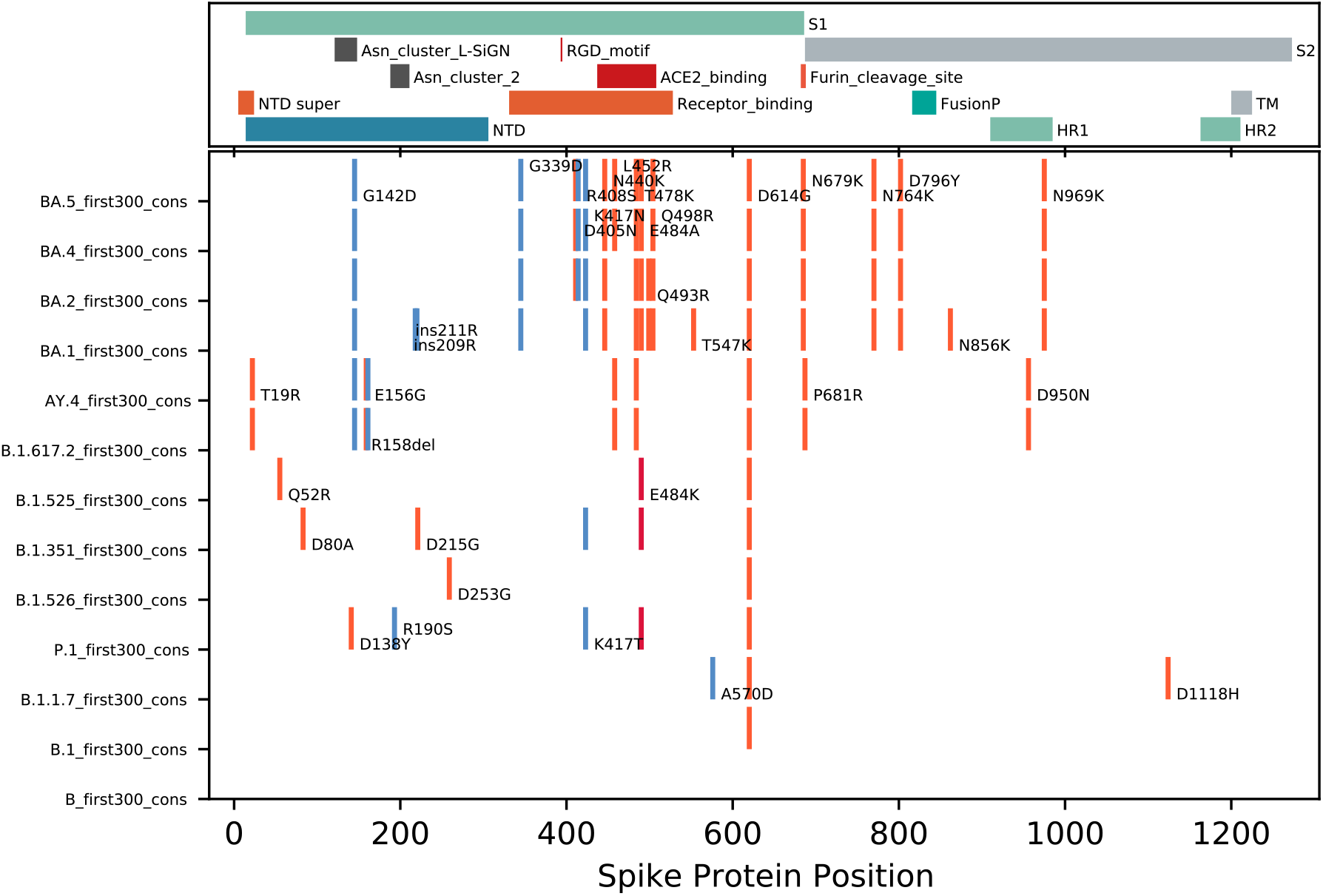
Location of charged amino acid changes in the spike protein. The spike protein sequences encoded by the first 300 reported genomes for the indicated SARS-CoV-2 lineages were collected, and charged amino acid changes from the original B lineage spike sequence were plotted. Charge changes were colored from dark blue (change from positive to negative charged amino acid (AA)), blue change from neutral to negative charged AA), orange (change from neutral to positive charged AA) and red (change from negative to positive charged AA). Substitutions are indicated by original AA/position in reference sequence spike/novel AA. The GenBank NC_045512 genome was used as reference. Key spike protein features of the SARS-CoV-2 spike protein are shown in the upper panel of the figure.

## Notes

### Competing Interest Statement

The authors have declared no competing interest.

### Summary of Updates

Analyses updated with data to mid-September 2022, New analysis (Figure 3) showing fraction of new genomes with changes from majority spike charge pattern (addressed the question: has the spike reached a maximum charge?).

